# Combinatorial *in silico* approach for cancer-associated 4Fe-4S protein discovery

**DOI:** 10.1101/2023.11.08.566327

**Authors:** Winnie W. L. Tam, Michael H. W. Cheng, Xiaoyong Mo, Bei-Bei He, Ulrike F. M. Ngo, Nicholas M. H. Yuen, Angela Y. L. Yau, Nicholas C. Wu, Edmund C. M. Tse

**Author notes:** **Corresponding Author** ECMT.

## Abstract

Iron-sulfur (Fe-S) proteins play vital roles in multiple cellular processes, including mediating redox balance as well as DNA replication and repair. Given the role of Fe-S cofactors in genome maintenance, mutations in such metalloproteins could be associated with cancer. Nevertheless, only a few cancer-associated Fe-S proteins have been identified. *In vitro*, Fe-S cluster is susceptible to degradation in oxic environment. It could also be replaced by other metal ions during protein purification, mis-labelled as zinc finger or Zn-containing proteins. *In silico*, bioinorganic Fe-S cluster lacks unique sequence characteristics that distinguish itself from other metal-coordination sites, making motif prediction based solely on protein sequence difficult. Thus, in this study, three traits have been employed to discover putative cancer-associated 4Fe-4S proteins. Here, we have analyzed the human proteome via a three-pronged approach: (i) the presence of a triamino acid motif, (ii) the geometric arrangements of the cysteines, and (iii) the mutations of cancer-associated cysteines. In addition to MUTYH, a known 4Fe-4S human DNA glycosylase, 21 novel proteins were discovered as potential cancer-associated 4Fe-4S proteins. While 6 receptor proteins and 3 growth factors have been identified as potential targets in this study, 5 histone lysine methyltransferases with SET domains were also predicted to contain 4Fe-4S metallocofactors. This work provides insights for rational adjustments in experimental design and novel cancer biomarker discovery.

## 1. Introduction

Iron-sulfur (Fe-S) clusters are essential in DNA replication and genome integrity maintainance.^1-5^ The coordination of Fe-S clusters is crucial for DNA replication and repair enzymes to carry out their physiological functions.^6^ DNA primase, an enzyme that initiates DNA replication by synthesizing short RNA primers complementary to DNA template, contains a 4Fe-4S cluster in proximity to DNA binding site (Figure 1A).^2^ Interference of Fe-S cluster binding pocket would affect the DNA binding and primer synthesis ability of DNA primase^1^. 4Fe-4S cluster could also be found in DNA replication helicase/nuclease 2 (DNA2) (Figure 1B), which is involved in Okazaki fragment maturation during DNA replication, replication checkpoint initiation, and telomere integrity maintenance^7-9^. Mutations of Fe-S cluster coordinating cysteine residues or residues adjacent to Fe-S domain in DNA2 could destabilize the Fe-S cluster, leading to decrease in nuclease activity and ATPase function^1^. Furthermore, DNA glycosylases EndoIII and MutY, which repair DNA damage via base excision repair (BER) mechanism, also contain Fe-S clusters in proximity to the DNA binding site (Figures 1C, 1D).^10-12^ Specifically, EndoIII removes oxidized pyrimidines while MutY removes adenine from A:8oxoG mispairs in genome^13^. The 4Fe-4S clusters in both EndoIII and MutY play a role in maintaining structural stability of the protein, while the cluster in EndoIII also facilitates localization of the DNA glycosylase at the point of DNA lesion by acting as an affinity switch^14-17^. As 4Fe-4S clusters are involved actively in DNA replication and repair, mutations in Fe-S protein may lead to genome instability and cancer formation.^18-19^

**Figure 1.**
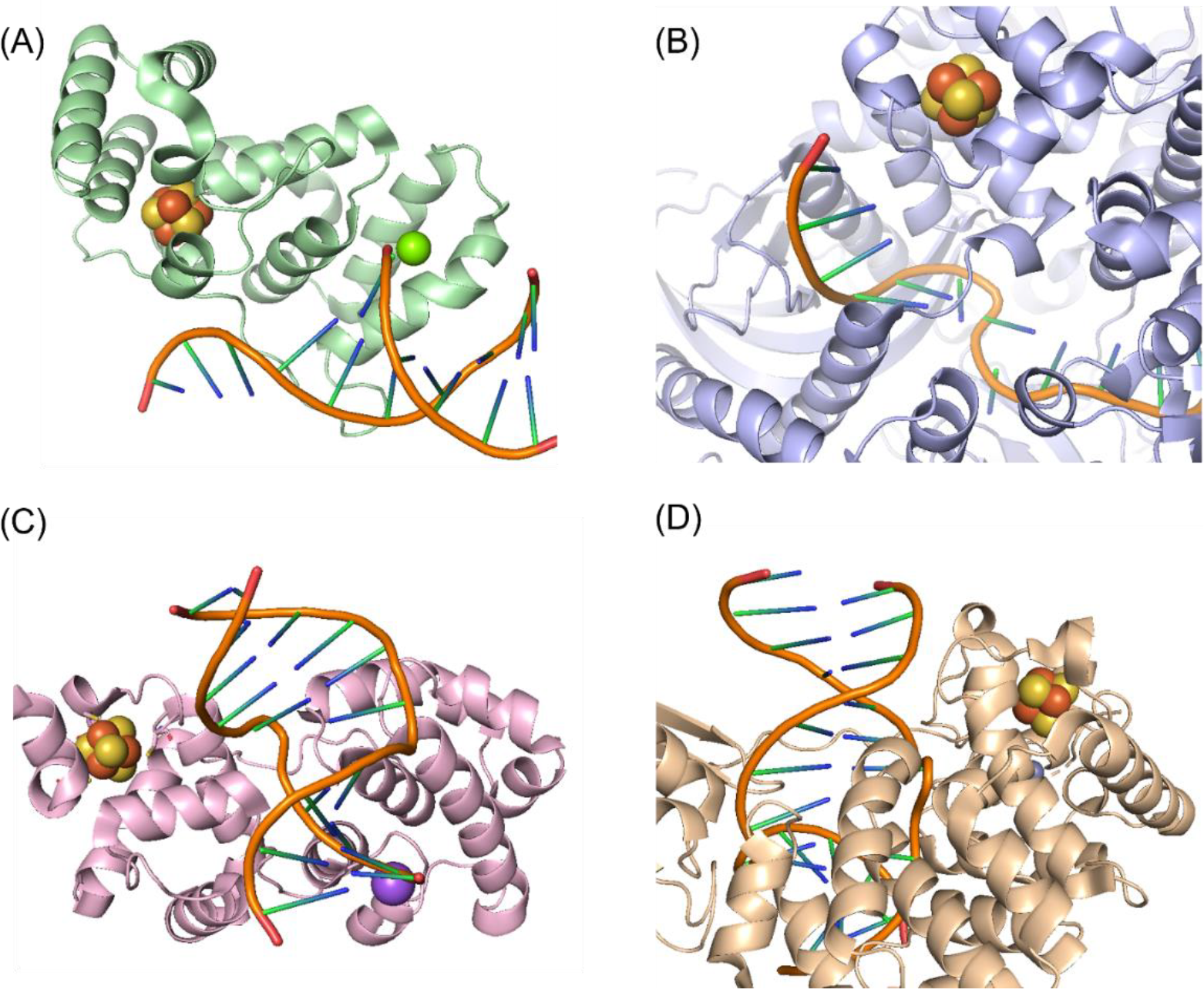
Fe-S cluster in (A) DNA primase, (B) DNA2, (C) EndoIII, and (D) MutY. PDB ID: 5F0Q, 5EAX, 1P59, and 7EF8. In all these crystal structures of DNA interacting protein, Fe-S clusters coordinate near the bound DNA.

For instance, mutations of cluster ligating cysteines in Fe-S protein were found to be associated with colon cancer and prostate cancer^20-22^. A cysteine-to-tryptophan substitution at position 306 (C306W) in MUTYH, the human analogue of MutY in *E. coli*, is associated with colonic polyposis and early onset of colon cancer^20^. Loss of cluster coordinating cysteines in MUTYH leads to destabilization of 4Fe-4S cluster, resulting in loss of DNA binding ability and enzyme activity^20, 23^. Likewise, Paulo et al. identified the C283R mutation in Fanconi anaemia group J protein (FANCJ, also known as BIRP1), a DNA helicase responsible for unwinding G-quadruplex (G4) DNA during DNA replication, as a “likely/potentially pathogenic” missense variant of prostate cancer by targeted next generation sequencing (NGS) panel^21^. However, these studies are limited to one specific Fe-S protein or a type of cancer, and there is no general profile of cancer-associated cysteine mutations in Fe-S proteins available currently.

Difficulty in characterizing Fe-S protein *in vitro* and *in silico* may account for the slow discovery of novel Fe-S protein. *In vitro*, like other metal centres, Fe-S clusters are sensitive to expression and aerobic purification conditions^24-26^. In particular, the clusters are prone to oxidative damage, which makes the isolation of holoprotein with intact clusters difficult^25^. The sensitivity of the clusters to oxygen varies as the electrostatic environment changes upon DNA binding^24^. Thus, Fe-S protein must be purified anaerobically in order to prevent oxidative degradation of the clusters^24-25^. Such stringent conditions hinder the isolation of Fe-S protein. *In silico*, predicting Fe-S binding site is challenging since there is no conserved sequence for 4Fe-4S cluster binding in high-potential Fe-S protein (HiPIPs), especially those involved in DNA repair and replication.^27^ In contrast to the high specificity of DNA-protein binding or protein-protein binding, 4Fe-4S cluster has a rather rudimentary structure that requires only 4 cysteine residues in sequence for coordination.^28^ There is a lack of strong predictor for the class of nucleic acid binding protein in linear model^29^. Some 4Fe-4S protein, such as XPD helicase and FANCJ, have a distinct 4Fe-4S domain while in some cases, like DNA2, the cluster coordinating cysteine residues were found to be distributed across the nuclease domain in an atypical spacing fashion.^1, 30-31^ It took up to 8 years after finding a mutation in cysteine being synthetically lethal in eukaryotic DNA polymerase to associate the mutation to a Fe-S cluster^32^. Developing robust methods to identify Fe-S protein in human proteome *in vitro* and *in silico* has been a major target in the fields of bioinorganic and biomedical chemistry.

Nevertheless, recent studies on Fe-S cluster assembly mechanism have unveiled new clues for identifying Fe-S protein.^33^ Atkinson et al. elucidated that the LYR motif is important for the insertion of Fe-S cluster into target protein in yeast mitochondria^34^. Later, Maio et al. identified that HSC20, a cochaperone for Fe-S cluster biogenesis in human, also recognises the L(I)YR motif and KKX_(6-10)_KK (X denotes proteinogenic amino acid) on succinate dehydrogenase B to facilitate the transfer of Fe-S cluster from ISCU-chaperone-cochaperone complex^35^. Similar motifs could also be found on other HSC20 interacting protein including SDHAF1, LYRM7, and GLRX5^35^. Mutations in such motifs was observed in various cancers^36-38^. These sequences could serve as new searching motifs for identification of novel 4Fe-4S protein in human.^39^

In this study, we propose a new algorithm to identify cancer-associated 4Fe-4S protein. Instead of limiting our search to known Fe-S protein, we tried to incorporate knowledge of Fe-S clusters biogenesis as well as chemical features of and structural constraints on known Fe-S protein to generate a list of putative 4Fe-4S cluster containing proteins. To confirm the presence of functional cysteines in such proteins, the list of putative 4Fe-4S protein was then compared with a database of cancer-associated somatic mutation to correlate cysteine mutated variants with human cancer. By identifying cancer-associated 4Fe-4S proteins, a new era in cancer prediction, prognosis, or treatment may be established.

## 2. Materials and Methods

Here we developed a combinatorial approach to aid identification of 4Fe-4S cluster containing protein. We analysed human proteome from multiple perspectives, the ability to receive 4Fe-4S cluster during biogenesis, potential Fe-S cluster binding site and its relation to cancer. These 3 bioinformatic approaches were combined to narrow down the searching space for novel 4Fe-4S proteins, which could be used as biomarkers for early detection and treatment targets. The workflow of our proposed algorithm is illustrated in Figure 2.

**Figure 2.**
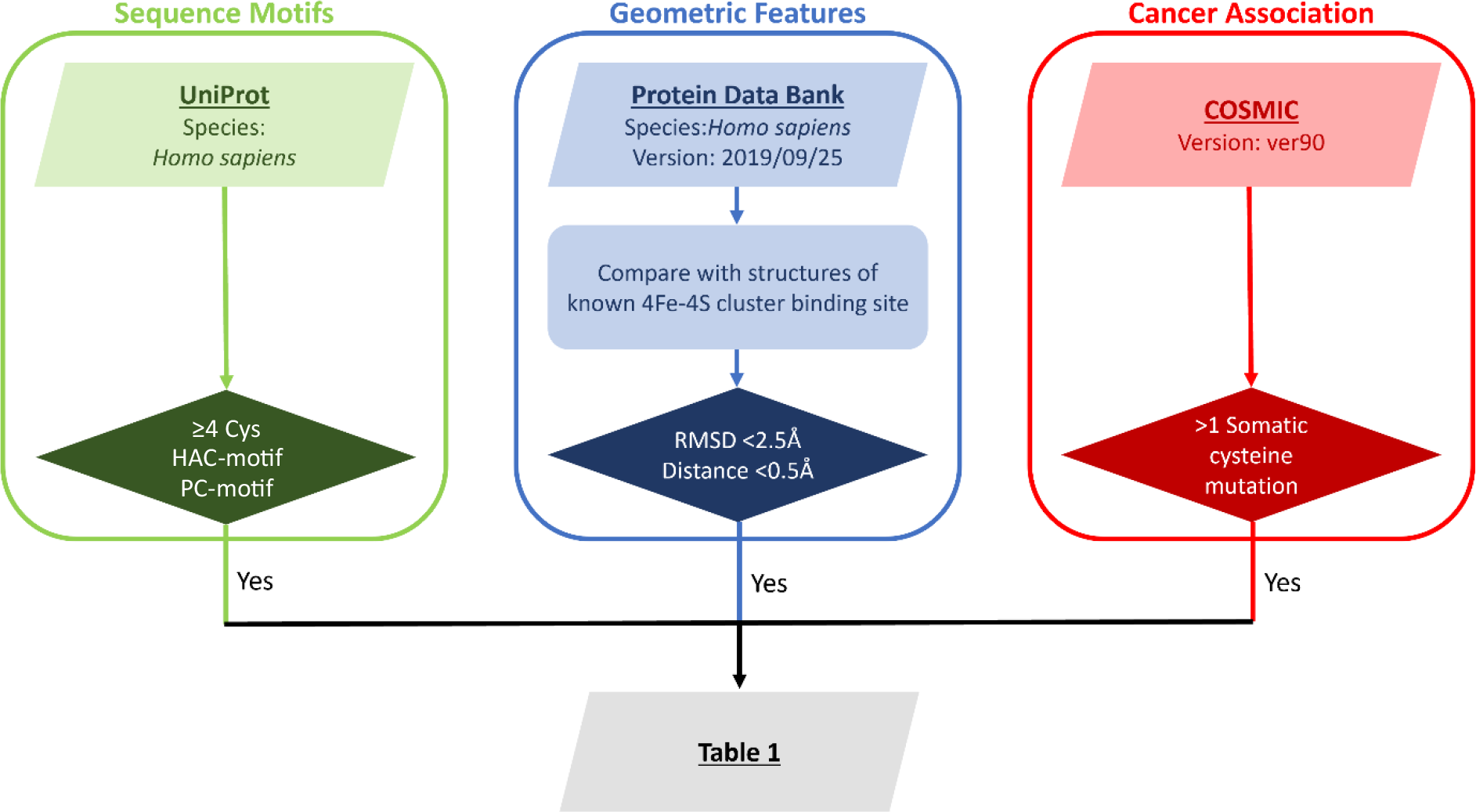
Workflow algorithm for identification of cancer-association 4Fe-4S cluster containing proteins. Protein entries from three databases were analysed in parallel. Final hits were overlapped and compiled manually.

### 2.1. Motif-based analysis

Human proteomes from UniProt were screened by two main criteria. First, the hit must contain at least four cysteine residues for 4Fe-4S cluster coordination. Second, the protein sequence must contain at least a set of Hydrophobic-Aromatic-Cationic (HAC) residues and two consecutive positively charged (PC) residues. In the HAC motif, hydrophobic residues could be valine, proline, methionine, alanine, isoleucine, or leucine; aromatic residues could be tyrosine or phenylalanine; cationic residues could be arginine or lysine. The pair of consecutive PC residues could be arginine or lysine, i.e. KK, RK, KR, or RR. Protein fulfilling both criteria would be counted as hits.

### 2.2. Cysteine geometry analysis

Jess, an algorithm for constrained-based structural template matching, was employed to evaluate if the geometry of cysteine in a protein could accommodate 4Fe-4S cluster^40^. Cysteine geometries were extracted from two known 4Fe-4S containing proteins (PDB ID: 5DII, 5MKQ) as templates (Figure 3A). Cysteine geometries of 6 other proteins (PDB ID: 1JQ4, 1QT9, 1ROE, 2DE7, 5MKO, 3QQ5), including 2Fe-2S binding or non-Fe-S binding, were extracted as negative templates (Figures 3B, 3C, 3D). These templates were used to search within the known human protein structure from the PDB data bank (PDB version: 2019.09.25). Best alignment with RMSD less than 2.5 Å and Cys-Cys distance between 5.8 Å and 6.8 Å as shown in Figure 3A, was counted as potential 4Fe-4S binding.

**Figure 3.**
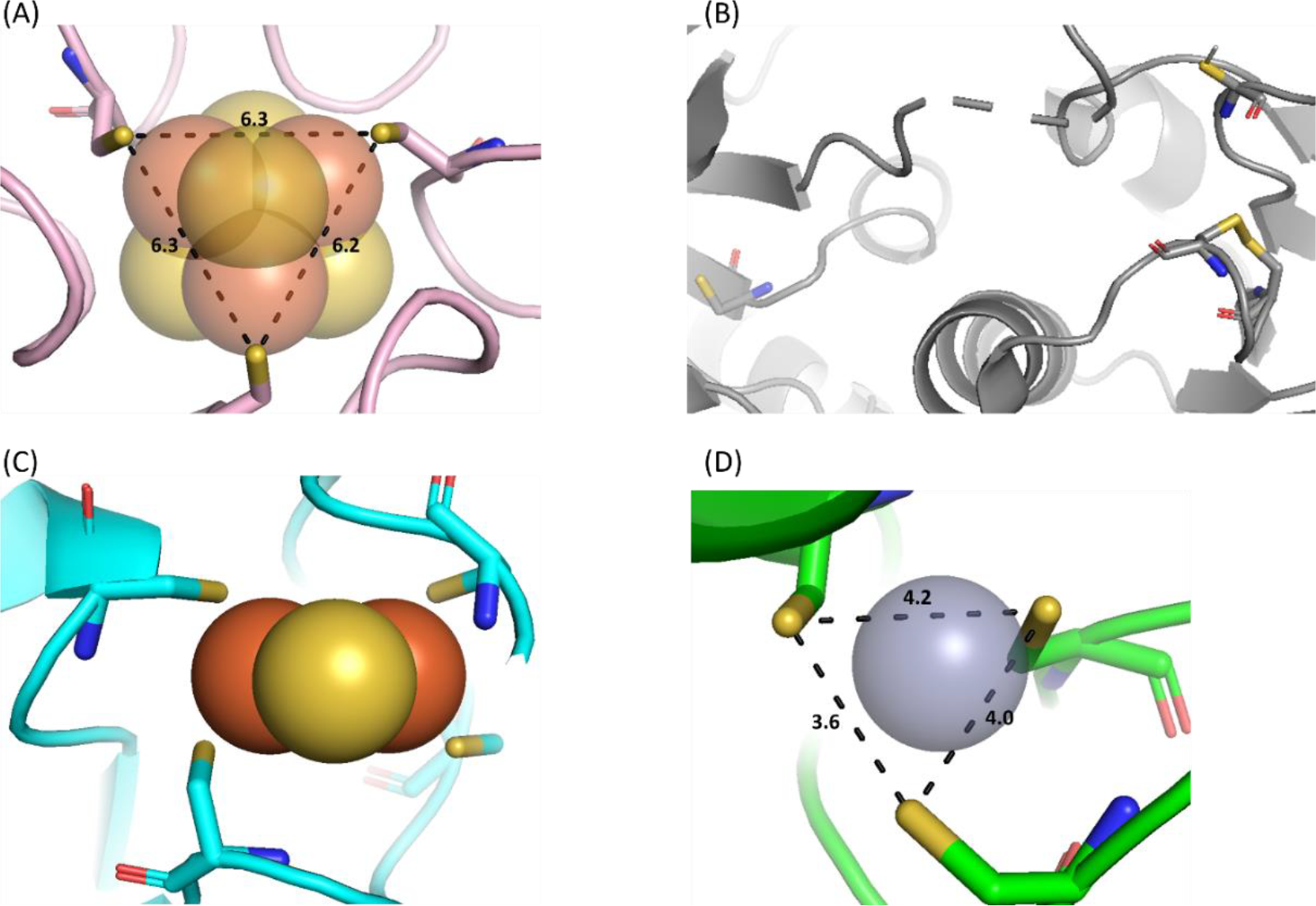
Structures served as template in cysteine geometry analysis. Distance between sulphur atoms of cysteine residues was measured in Å. (A) 4Fe-4S binding site of microcompartment shell protein (PDB ID: 5DII). Three sulphur atoms of the coordinating cysteine form an equilateral triangle of around 6 Å on each side. (B) Geometries of cysteine in non-Fe-S binding site as negative template (PDB ID: 3QQ5). (C) 2Fe-2S binding site of ferredoxin (PDB ID: 1QT9) as negative template. (D) Zn binding site of TtuA (PDB ID: 5MKO) as negative template.

### 2.3. Binding site mutations in cancer

Protein with at least one somatic cysteine mutation reported in cancer was extracted from Catalogue of Somatic Mutations in Cancer (COSMIC) database ver90. The outcomes of the selection processes were overlaid manually. A more detailed analysis on structure, homology, and protein function was done on hits fulfilling all criteria.

## 3. Results and discussion

### 3.1. Potential cancer-associated 4Fe-4S cluster containing protein identified in our study

The number of hits fulfilling each criterion was shown in Figure 4. Out of 73101 human proteins in UniProt, 22019 hits were identified as potential Fe-S cluster recipient proteins because they contain at least 4 cysteine residues, HAC motif, and PC motif. Since it was reported that both leucine and isoleucine at the first position of the LYR motif could be recognized by the ISCU chaperone-cochaperone complex, here we expanded the search into HAC motif to allow substitution of amino acids with similar properties^35^. From investigating the geometry of cysteine, in particular the sulfur atom, we were able to identify 124 targets from 16926 known human protein structures. Out of 74349 entries in the COSMIC database, 11841 proteins with at least one somatic cysteine mutation were reported in human cancer. Listed in Table 1, 22 proteins fulfilling all three criteria were considered as potential cancer-associated 4Fe-4S cluster containing proteins. The algorithm has successfully recalled the known cancer-associated 4Fe-4S cluster containing protein, MUTYH. On top of that, a wide range of proteins including methyltransferase, growth factor, and receptor has been identified as potentially 4Fe-4S binding. The cysteine residues in most of these proteins were previously identified as Tri-/Bi-Zn binding or containing extensive disulphide bridges.

**Figure 4.**
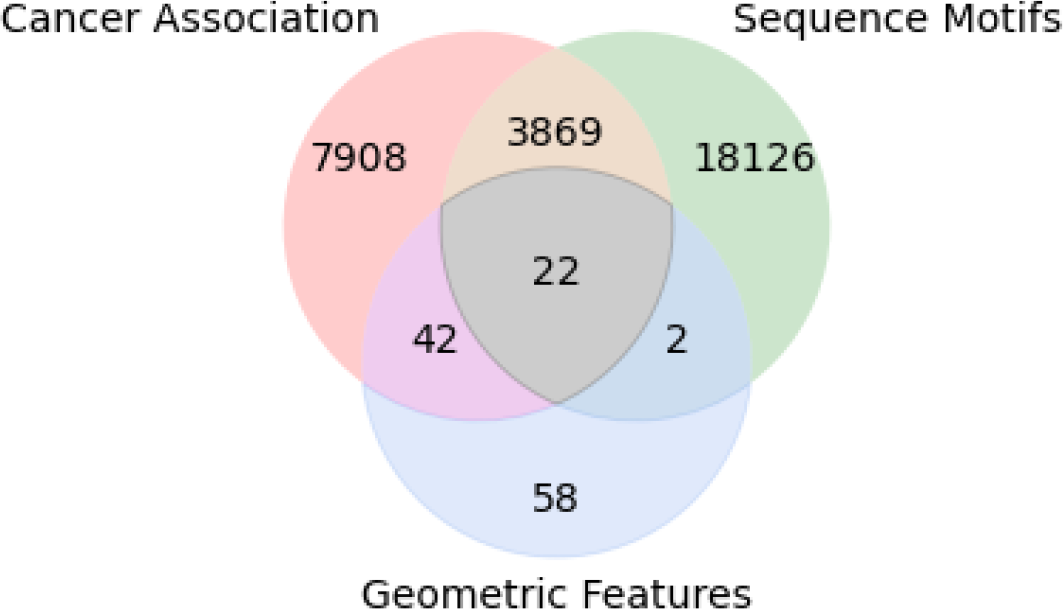
Overlap of results from 3 databases. 22 hits fulfilling all 3 criteria were listed in Table 1.

**Table 1.**
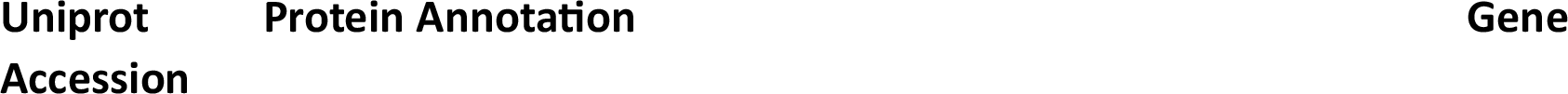
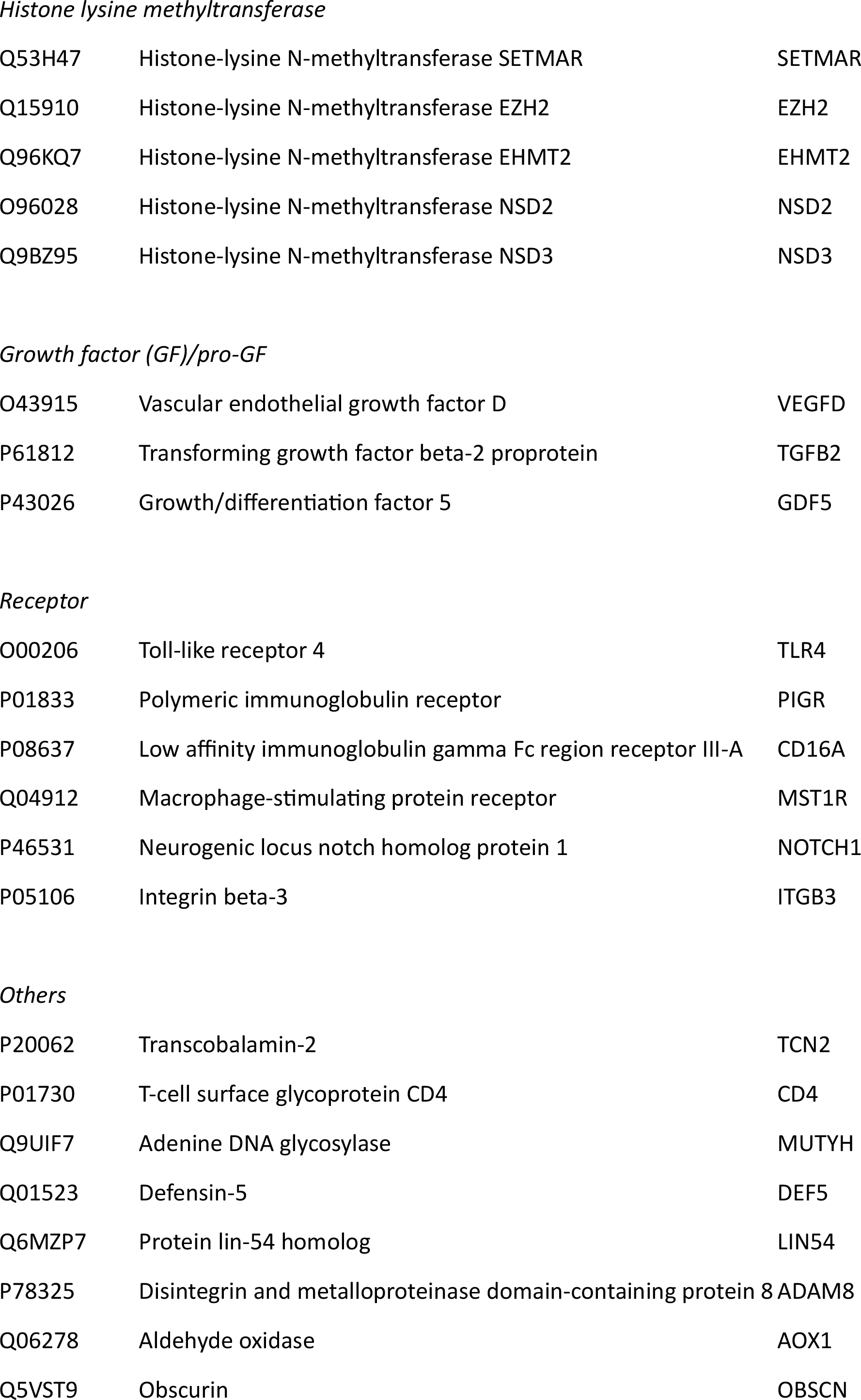
Lists of proteins identified as putative cancer-associated 4Fe-4S cluster containing proteins. Proteins were grouped according to their classes.

### 3.2. Histone lysine N-methyltransferase as potential Fe-S protein

In our lists of hits, 4 of them (EZH2, EHMT2, NSD2, NSD3) belong to class V-like SAM-binding methyltransferase superfamily histone-lysine methyltransferase family. Though SETMAR, a fusion of methylase and transposase, does not belong to histone-lysine methyltransferase family, it methylates histone protein using *S*-adenosyl-methionine (SAM) as substrate like other histone-lysine methyltransferases. Sequence and structural analysis were performed to further investigate and annotate possible 4Fe-4S binding site. Domain distribution of these protein was predicted by PROSITE and illustrated in Figure 5A^41^. All proteins contain a cysteine rich region *N*-terminal to the SET domain. Though no significant sequence similarity was found in pre-SET and AWS domain, superposition of crystal structure suggests structural similarity between the pre-SET and AWS domains (Figure 5B). Zoom in for the potential 4Fe-4S binding site was shown in Figure 5C. Other proteins in this family do not possess such domain, and thus, were not included in the final hits. EZH1, the closely related homolog of EZH2, was not included in the final hits due to the lack of 3D structural information at the time of search.

**Figure 5.**
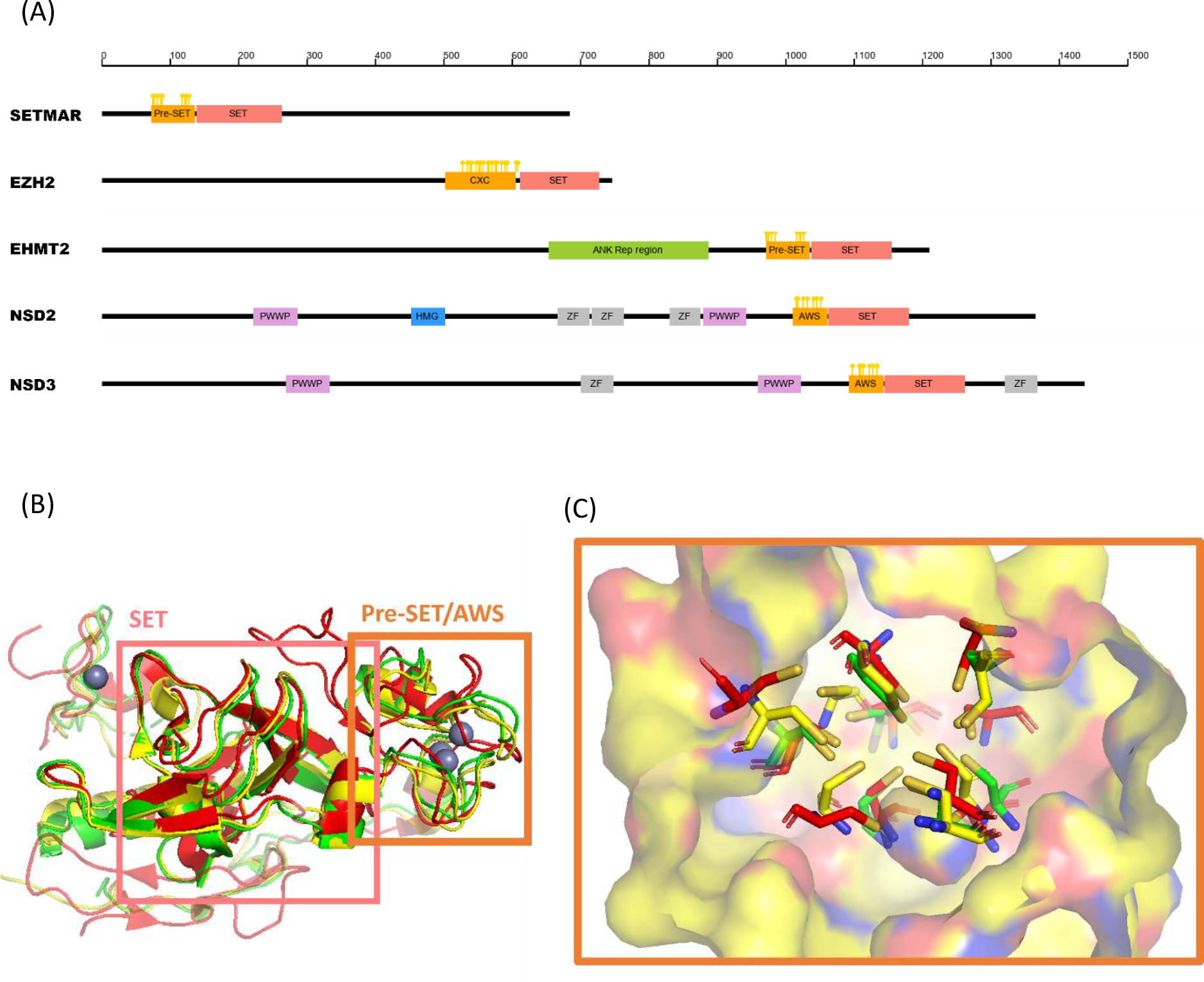
(A) Domain distribution of histone lysine methyltransferase. Cysteine residues in cysteine-rich regions were labelled by yellow mark. The predicted 4Fe-4S binding sites were *N*-terminal to SET domain. (B) Superimposed structures of EHMT2 (PDB ID: 2O8J), NSD2 (PDB ID: 5LSU), and NSD3 (PDB ID: 5UPD). Pre-SET and AWS domains are similar in structure, and each contains a tri-zinc binding site. (C) Zoom in of the tri-zinc binding site. 9 invariant cysteine residues form a pocket for coordination of three zinc ions in a triangular cluster. It was predicted that this pocket may coordinate 4Fe-4S cluster instead of Zn *in vivo*.

Pre-SET domain is characterised by a cysteine-rich region which coordinates 3 Zn ions. 9 cysteine residues are grouped in two protein segments, separated by a long random coil with variable length^42^. This *N*-terminal flanking cysteine-rich region is common in SUV39-family protein including EHMT2. Although pre-SET does not contribute to the catalytic activity of methyltransferase, it was shown that both *N*-terminal and *C*-terminal adjacent cysteine-rich regions were essential for H3-specific histone methyltransferases (HMTase) activity^43^. X-ray crystallography structure showed extensive interactions between pre-SET domain and SET domain, implying the role of pre-SET in stabilizing the structure of SET and the extended binding site for the histone tail^44^. In other SAM-dependent enzymes in human, such as molybdenum cofactor biosynthesis protein 1 and viperin, 4Fe-4S catalyses the formation of SAM radicals that lead to the reductive cleavage of SAM^45-46^. Methyltransferases in bacterial system were found to utilize the 4Fe-4S-SAM radical pathway to methylate RNA^47^. Our study identified histone lysine methyltransferases as a novel class of protein that potentially coordinates 4Fe-4S metallocofactor. *In vitro* study is required to validate the presence of 4Fe-4S cluster.

Histone lysine methyltransferases play an important role in transcription regulation, and their mutatants could lead to dysregulation of epigenetic regulation, a feature in cancer cell^48^. Methylations at H3K9 and H3K27 are usually associated with gene silencing while H3K4 and H3K36 are usually associated with gene activity. Bivalent methylation at H3K4 and H3K27 was observed in early embryogenesis to maintain pluripotency of embryonic stem cells^49^. Different epigenetic profiles between cancer cell and normal cell suggest histone methylation landscape are associated with formation and progression of cancer^50^.

### 3.3. Aldehyde oxidase

Only two hits on our list have been previously annotated as Fe-S binding, MUTYH and aldehyde oxidase (AOX1). While it is known that MUTYH binds a 4Fe-4S cluster, X-ray crystallographic information of AOX1 revealed that it contains two 2Fe-2S clusters in proximity (Figure 6). 4Fe-4S cluster is oxygen reactive and may be converted to 2Fe-2S cluster under aerobic condition^51^. Such oxygen switch has been employed in fumerate and nitrate reduction (FNR) regulator to modulate expression of gene for anaerobic respiration^52-53^. Since the crystal obtained was under aerobic condition, it could be possible that AOX1 is 4Fe-4S binding under anaerobic condition.

**Figure 6.**
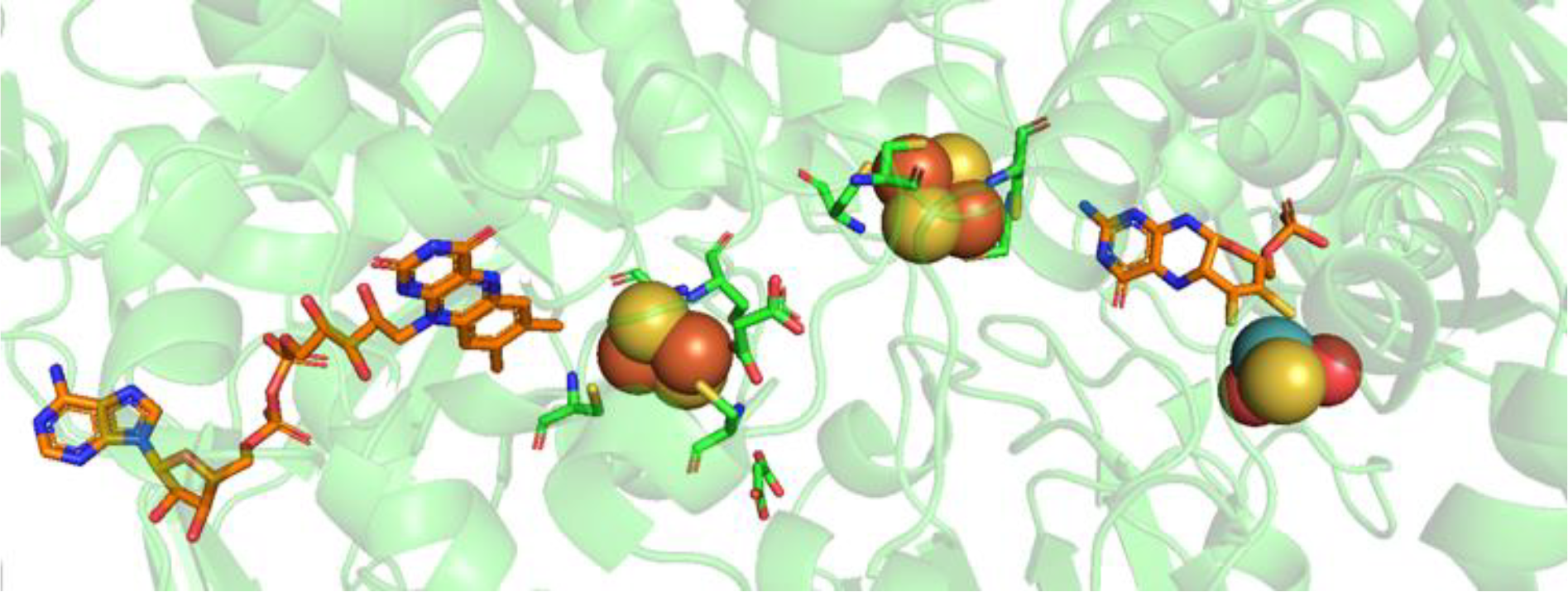
Crystal structure of AOX1 (PDB ID: 4UHW). Two 2Fe-2S clusters lie between FAD and the catalytic molybdenum cofactor.

As an important enzyme for detoxification of aldehydes and azaheterocycles, AOX1 was downregulated in various cancer^54-56^. Loss of AOX1 expression results in elevated cell proliferation and altered metabolic pathways including glucose metabolism and nucleotides synthesis^54^. On top of that, loss of AOX1 expression induces epithelial to mesenchymal transition, which results in cancer cell migration and invasion^54^. Thus, AOX1 could potentially be a therapeutic target in cancer.

## Conclusions

Here we unveiled novel putative 4Fe-4S cluster containing proteins associated with cancer. Though a vast number of metal binding site prediction tools are available, most of them fail to distinguish between Zn binding site and Fe-S binding site due to the similarity in the conformation of cysteine residues in both binding sites. Taking into account of the Fe-S cluster biogenesis pathway, this combinatorial approach drastically abridges the table of hits from three distinct and independent methods. Our results suggest an intriguing possibility of 4Fe-4S binding site in histone lysine methyltransferases as well as other receptors and growth factors. This work provides insights for further *in vitro* studies design, such as anaerobic protein purification, to verify the presence of 4Fe-4S clusters in metalloenzymes. Better understanding of these cancer-associated proteins would be beneficial to cancer prognosis and treatment.

## Acknowledgements

We acknowledge the funding support from “Laboratory for Synthetic Chemistry and Chemical Biology” under the Health@InnoHK Program launched by Innovation and Technology Commission, the Government of Hong Kong Special Administrative Region of the People’s Republic of China. E.C.M.T. would like to recognize the support from the URC Seed Fund for Basic Research, the RGC Research Matching Grant Schemes (RMGS: 207301212, 207301251), and the Croucher Foundation Start-up Allowance. N.C.W. would like to thank the funding from the Searle Scholars Program. We also express our gratitude to Nicholas H. M. Tam, Angela T. H. Fan, Claudia K. Y. Hui, Alice S. H. Yau, Carl Y. N. Cheung, Kyungmin Nam, Jacqueline Wu, Mary J. S. Yuen, and K. Ming Tam for their efforts in the *in-silico* screening process during their summer internships. We acknowledge the source code of Jess from Dr. Roman Laskowski and Dr. Ioannis Riziotis.

## References

(1) Paul, V. D.; Lill, R., Biogenesis of cytosolic and nuclear iron-sulfur proteins and their role in genome stability. Biochim. Biophys. Acta 2015, 1853, 1528–1539.

(2) Weiner, B. E.; Huang, H.; Dattilo, B. M.; Nilges, M. J.; Fanning, E.; Chazin, W. J., An iron-sulfur cluster in the C-terminal domain of the p58 subunit of human DNA primase. J. Biol. Chem. 2007, 282, 33444–33451.

(3) White, M. F.; Dillingham, M. S., Iron-sulphur clusters in nucleic acid processing enzymes. Curr. Opin. Struct. Biol. 2012, 22, 94–100.

(4) Maio, N.; Lafont, B. A. P.; Sil, D.; Li, Y.; Bollinger, J. M.; Krebs, C.; Pierson, T. C.; Linehan, W. M.; Rouault, T. A., Fe-S cofactors in the SARS-CoV-2 RNA-dependent RNA polymerase are potential antiviral targets. Science 2021, 373, 236–241.

(5) Py, B.; Barras, F., Building Fe–S proteins: bacterial strategies. Nat. Rev. Microbiol 2010, 8, 436–446.

(6) Lill, R., Function and biogenesis of iron–sulphur proteins. Nature 2009, 460, 831–838.

(7) Chen, X.; Niu, H.; Chung, W. H.; Zhu, Z.; Papusha, A.; Shim, E. Y.; Lee, S. E.; Sung, P.; Ira, G., Cell cycle regulation of DNA double-strand break end resection by Cdk1-dependent Dna2 phosphorylation. Nat. Struct. Mol. Biol. 2011, 18, 1015–1019.

(8) Kumar, S.; Burgers, P. M., Lagging strand maturation factor Dna2 is a component of the replication checkpoint initiation machinery. Genes Dev. 2013, 27, 313–321.

(9) Lin, W.; Sampathi, S.; Dai, H.; Liu, C.; Zhou, M.; Hu, J.; Huang, Q.; Campbell, J.; Shin-Ya, K.; Zheng, L.; Chai, W.; Shen, B., Mammalian DNA2 helicase/nuclease cleaves G-quadruplex DNA and is required for telomere integrity. EMBO J. 2013, 32, 1425–1439.

(10) Doetsch, P., Mechanisms of DNA Repair. Elsevier Science & Technology: San Diego, UNITED STATES, 2012.

(11) Zwang, T. J.; Tse, E. C. M.; Barton, J. K., Sensing DNA through DNA Charge Transport. ACS Chem. Biol. 2018, 13, 1799–1809.

(12) Tse, E. C. M.; Zwang, T. J.; Barton, J. K., The Oxidation State of [4Fe4S] Clusters Modulates the DNA-Binding Affinity of DNA Repair Proteins. J. Am. Chem. Soc. 2017, 139, 12784–12792.

(13) David, S. S.; Williams, S. D., Chemistry of Glycosylases and Endonucleases Involved in Base-Excision Repair. Chem. Rev. 1998, 98, 1221–1262.

(14) Rouault, T.; Ichiye, T.; Bonomi, F.; Booker, S.; Chakrabarti, M.; Einsle, O.; Fontecilla, J.; Hendrich, M.; Hille, R.; Hu, Y., Characterization, Properties and Applications. De Gruyter, Inc.: Berlin/Boston, GERMANY, 2017.

(15) O’Brien, E.; Holt, M. E.; Thompson, M. K.; Salay, L. E.; Ehlinger, A. C.; Chazin, W. J.; Barton, J. K., The [4Fe4S] cluster of human DNA primase functions as a redox switch using DNA charge transport. Science 2017, 355, eaag1789.

(16) Grodick, M. A.; Muren, N. B.; Barton, J. K., DNA Charge Transport within the Cell. Biochem. 2015, 54, 962–973.

(17) Tse, E. C. M.; Zwang, T. J.; Bedoya, S.; Barton, J. K., Effective Distance for DNA-Mediated Charge Transport between Repair Proteins. ACS Cent. Sci. 2019, 5, 65–72.

(18) Rouault, T. A., Mitochondrial iron overload: causes and consequences. Curr. Opin. Genet. Dev. 2016, 38, 31–37.

(19) Barton, J. K.; Silva, R. M. B.; O’Brien, E., Redox Chemistry in the Genome: Emergence of the [4Fe4S] Cofactor in Repair and Replication. Annu. Rev. Biochem. 2019, 88, 163–190.

(20) McDonnell, K. J.; Chemler, J. A.; Bartels, P. L.; O’Brien, E.; Marvin, M. L.; Ortega, J.; Stern, R. H.; Raskin, L.; Li, G. M.; Sherman, D. H.; Barton, J. K.; Gruber, S. B., A human MUTYH variant linking colonic polyposis to redox degradation of the [4Fe4S](2+) cluster. Nat. Chem. 2018, 10, 873–880.

(21) Paulo, P.; Maia, S.; Pinto, C.; Pinto, P.; Monteiro, A.; Peixoto, A.; Teixeira, M. R., Targeted next generation sequencing identifies functionally deleterious germline mutations in novel genes in early-onset/familial prostate cancer. PLoS Genet. 2018, 14, e1007355.

(22) Saxena, N.; Maio, N.; Crooks, D. R.; Ricketts, C. J.; Yang, Y.; Wei, M. H.; Fan, T. W.; Lane, A. N.; Sourbier, C.; Singh, A.; Killian, J. K.; Meltzer, P. S.; Vocke, C. D.; Rouault, T. A.; Linehan, W. M., SDHB-Deficient Cancers: The Role of Mutations That Impair Iron Sulfur Cluster Delivery. J. Natl. Cancer Inst. 2016, 108, djv287.

(23) Fontecave, M.; Ollagnier-de-Choudens, S., Iron–sulfur cluster biosynthesis in bacteria: Mechanisms of cluster assembly and transfer. Arch. Biochem. Biophys. 2008, 474, 226–237.

(24) David, S. S., Fe-S Cluster Enzymes Part B. Elsevier Science & Technology: San Diego, UNITED STATES, 2018.

(25) Rouault, T. A., Iron-sulfur proteins hiding in plain sight. Na.t Chem. Biol. 2015, 11, 442–445.

(26) Cvetkovic, A.; Menon, A. L.; Thorgersen, M. P.; Scott, J. W.; Poole, F. L., 2nd; Jenney, F. E., Jr.; Lancaster, W. A.; Praissman, J. L.; Shanmukh, S.; Vaccaro, B. J.; Trauger, S. A.; Kalisiak, E.; Apon, J. V.; Siuzdak, G.; Yannone, S. M.; Tainer, J. A.; Adams, M. W., Microbial metalloproteomes are largely uncharacterized. Nature 2010, 466, 779–782.

(27) Fuss, J. O.; Tsai, C. L.; Ishida, J. P.; Tainer, J. A., Emerging critical roles of Fe-S clusters in DNA replication and repair. Biochim. Biophys. Acta 2015, 1853, 1253–1271.

(28) Lambert, A. R.; Sussman, D.; Shen, B.; Maunus, R.; Nix, J.; Samuelson, J.; Xu, S.-Y.; Stoddard, B. L., Structures of the Rare-Cutting Restriction Endonuclease NotI Reveal a Unique Metal Binding Fold Involved in DNA Binding. Struct. 2008, 16, 558–569.

(29) Estellon, J.; Ollagnier de Choudens, S.; Smadja, M.; Fontecave, M.; Vandenbrouck, Y., An integrative computational model for large-scale identification of metalloproteins in microbial genomes: a focus on iron-sulfur cluster proteins. Metallomics 2014, 6, 1913–1930.

(30) Wu, Y.; Brosh, R. M., Jr., DNA helicase and helicase-nuclease enzymes with a conserved iron-sulfur cluster. Nucleic Acids Res. 2012, 40, 4247–4260.

(31) Netz, D. J. A.; Stith, C. M.; Stümpfig, M.; Köpf, G.; Vogel, D.; Genau, H. M.; Stodola, J. L.; Lill, R.; Burgers, P. M. J.; Pierik, A. J., Eukaryotic DNA polymerases require an iron-sulfur cluster for the formation of active complexes. Nat. Chem. Biol. 2012, 8, 125–132.

(32) Baranovskiy, A. G.; Siebler, H. M.; Pavlov, Y. I.; Tahirov, T. H., Iron-Sulfur Clusters in DNA Polymerases and Primases of Eukaryotes. Meth. Enzymol. 2018, 599, 1–20.

(33) Maio, N.; Rouault, T. A., Iron–sulfur cluster biogenesis in mammalian cells: New insights into the molecular mechanisms of cluster delivery. Biochim. Biophys. Acta 2015, 1853, 1493–1512.

(34) Atkinson, A.; Smith, P.; Fox, J. L.; Cui, T. Z.; Khalimonchuk, O.; Winge, D. R., The LYR protein Mzm1 functions in the insertion of the Rieske Fe/S protein in yeast mitochondria. Mol. Cell. Biol. 2011, 31, 3988–3996.

(35) Maio, N.; Singh, A.; Uhrigshardt, H.; Saxena, N.; Tong, W. H.; Rouault, T. A., Cochaperone binding to LYR motifs confers specificity of iron sulfur cluster delivery. Cell. Metab. 2014, 19, 445–457.

(36) Amar, L.; Bertherat, J.; Baudin, E.; Ajzenberg, C.; Bressac-de Paillerets, B.; Chabre, O.; Chamontin, B.; Delemer, B.; Giraud, S.; Murat, A.; Niccoli-Sire, P.; Richard, S.; Rohmer, V.; Sadoul, J. L.; Strompf, L.; Schlumberger, M.; Bertagna, X.; Plouin, P. F.; Jeunemaitre, X.; Gimenez-Roqueplo, A. P., Genetic testing in pheochromocytoma or functional paraganglioma. J. Clin. Oncol. 2005, 23, 8812–8818.

(37) Celestino, R.; Lima, J.; Faustino, A.; Maximo, V.; Gouveia, A.; Vinagre, J.; Soares, P.; Lopes, J. M., A novel germline SDHB mutation in a gastrointestinal stromal tumor patient without bona fide features of the Carney-Stratakis dyad. Fam. Cancer 2012, 11, 189–194.

(38) Ricketts, C. J.; Shuch, B.; Vocke, C. D.; Metwalli, A. R.; Bratslavsky, G.; Middelton, L.; Yang, Y.; Wei, M. H.; Pautler, S. E.; Peterson, J.; Stolle, C. A.; Zbar, B.; Merino, M. J.; Schmidt, L. S.; Pinto, P. A.; Srinivasan, R.; Pacak, K.; Linehan, W. M., Succinate dehydrogenase kidney cancer: an aggressive example of the Warburg effect in cancer. J. Urol. 2012, 188, 2063–2071.

(39) Rouault, T. A., Biogenesis of iron-sulfur clusters in mammalian cells: new insights and relevance to human disease. Dis. model. mech. 2012, 5, 155–164.

(40) Barker, J. A.; Thornton, J. M., An algorithm for constraint-based structural template matching: application to 3D templates with statistical analysis. Bioinformatics 2003, 19, 1644–1649.

(41) de Castro, E.; Sigrist, C. J.; Gattiker, A.; Bulliard, V.; Langendijk-Genevaux, P. S.; Gasteiger, E.; Bairoch, A.; Hulo, N., ScanProsite: detection of PROSITE signature matches and ProRule-associated functional and structural residues in proteins. Nucleic Acids Res. 2006, 34, W362–W365.

(42) Dillon, S. C.; Zhang, X.; Trievel, R. C.; Cheng, X., The SET-domain protein superfamily: protein lysine methyltransferases. Genome Biol. 2005, 6, 227.

(43) Rea, S.; Eisenhaber, F.; O’Carroll, D.; Strahl, B. D.; Sun, Z. W.; Schmid, M.; Opravil, S.; Mechtler, K.; Ponting, C. P.; Allis, C. D.; Jenuwein, T., Regulation of chromatin structure by site-specific histone H3 methyltransferases. Nature 2000, 406, 593–599.

(44) Wilson, J. R.; Jing, C.; Walker, P. A.; Martin, S. R.; Howell, S. A.; Blackburn, G. M.; Gamblin, S. J.; Xiao, B., Crystal structure and functional analysis of the histone methyltransferase SET7/9. Cell 2002, 111, 105–115.

(45) Duschene, K. S.; Broderick, J. B., The antiviral protein viperin is a radical SAM enzyme. FEBS Lett. 2010, 584, 1263–1267.

(46) Hanzelmann, P.; Hernandez, H. L.; Menzel, C.; Garcia-Serres, R.; Huynh, B. H.; Johnson, M. K.; Mendel, R. R.; Schindelin, H., Characterization of MOCS1A, an oxygen-sensitive iron-sulfur protein involved in human molybdenum cofactor biosynthesis. J. Biol. Chem. 2004, 279, 34721–34732.

(47) Allen, K. D.; Wang, S. C., Initial characterization of Fom3 from Streptomyces wedmorensis: The methyltransferase in fosfomycin biosynthesis. Arch. Biochem. Biophys. 2014, 543, 67–73.

(48) Muntean, A. G.; Hess, J. L., Epigenetic dysregulation in cancer. Am. J. Pathol. 2009, 175, 1353–1361.

(49) Vastenhouw, N. L.; Schier, A. F., Bivalent histone modifications in early embryogenesis. Curr. Opin. Cell Biol. 2012, 24, 374–386.

(50) Varier, R. A.; Timmers, H. T., Histone lysine methylation and demethylation pathways in cancer. Biochim. Biophys. Acta 2011, 1815, 75–89.

(51) Oakley, K. M.; Lehane, R. L.; Zhao, Z.; Kim, E., Dioxygen reactivity of a biomimetic [4Fe-4S] compound exhibits [4Fe-4S] to [2Fe-2S] cluster conversion. J. Inorg. Biochem. 2022, 228, 111714.

(52) Crack, J. C.; Green, J.; Thomson, A. J.; Le Brun, N. E., Iron-sulfur clusters as biological sensors: the chemistry of reactions with molecular oxygen and nitric oxide. Acc. Chem. Res. 2014, 47, 3196–3205.

(53) Khoroshilova, N.; Popescu, C.; Munck, E.; Beinert, H.; Kiley, P. J., Iron-sulfur cluster disassembly in the FNR protein of Escherichia coli by O2: [4Fe-4S] to [2Fe-2S] conversion with loss of biological activity. Proc. Natl. Acad. Sci. U.S.A. 1997, 94, 6087–6092.

(54) Vantaku, V.; Putluri, V.; Bader, D. A.; Maity, S.; Ma, J.; Arnold, J. M.; Rajapakshe, K.; Donepudi, S. R.; von Rundstedt, F. C.; Devarakonda, V.; Dubrulle, J.; Karanam, B.; McGuire, S. E.; Stossi, F.; Jain, A. K.; Coarfa, C.; Cao, Q.; Sikora, A. G.; Villanueva, H.; Kavuri, S. M., et al., Epigenetic loss of AOX1 expression via EZH2 leads to metabolic deregulations and promotes bladder cancer progression. Oncogene 2020, 39, 6265–6285.

(55) Xiong, L.; Feng, Y.; Hu, W.; Tan, J.; Li, S.; Wang, H., Expression of AOX1 Predicts Prognosis of Clear Cell Renal Cell Carcinoma. Front. genet. 2021, 12, 683173.

(56) Zhang, W.; Chai, W.; Zhu, Z.; Li, X., Aldehyde oxidase 1 promoted the occurrence and development of colorectal cancer by up-regulation of expression of CD133. Int. Immunopharmacol. 2020, 85, 106618.

